# Mimicry of dopamine 1 receptor signaling with Cell-Penetrating Peptides

**DOI:** 10.1101/2020.01.07.897470

**Authors:** Nicola Lorenzon, Maxime Gestin, Ülo Langel

## Abstract

In this study, through the use of protein mimicry, a peptide was developed to activate the dopamine 1 receptor signaling pathway in substitution of L-DOPA. The peptide proved to be capable of efficiently ubiquitously penetrating the cell without the need for transfection agents or chiral recognition by specific pathways. Furthermore, the peptide induced the cellular response normally achieved through the activation of the receptor in cells that had not been treated with the natural ligand. The peptide could work as a candidate substitute to L-DOPA, leading the way for a peptides-based treatment for Parkinson’s disease.

## Introduction

Cell-penetrating peptides (CPPs) are a class of short generally cationic and/or amphipathic peptides capable of internalizing the cell via different mechanisms (Bechara and Sagan 2013). This property is bestowed by the presence of prevalently positively-charged and hydrophobic amino acid residues in the sequence, which interact with the negatively-charged components of the lipidic bilayer of the cell membrane (Madani et al. 2011).

The use of CPPs to transfect cargoes (such as small molecules, small proteins, peptides or nucleic acids) into the cell has been widely established (Li et al. 2015; Singh et al. 2018; Kurrikoff and Langel 2019). As vectors, CPPs can be used either in complex with the molecules of interest (e.g. forming electrostatically-bound particles that dissociate once transfected) or be covalently conjugated to them (Gagat et al. 2017). In particular, by conjugating lipidated CPPs with an amino acid sequence resembling parts of a functional full-length protein, chimeric peptides, generally labeled as ‘pepducins’, can be produced. Thanks to these functional domains on their backbone, pepducins are capable, once internalized, of specifically interfering with protein-protein interactions by competitively binding to the target protein (Zhang et al. 2015). This has been used in the past, for example, to modulate the response of tyrosine-kinase and angiotensin receptors and clinical trials have recently started for an antiplatelet-agent-acting pepducin (Yu et al. 2010; Yu et al. 2015; Gurbel et al. 2016). Furthermore, pepducins represent a promising class of allosteric G-protein coupled receptors (GPCRs) modulators, overcoming the selectivity problem associated with the use of extracellular ligands (Eglen and Reisine 2011). For example, one neurotransmitter (NT) is often capable of inducing multiple and opposite responses depending on the receptor it binds to, which is why the specific modulation of a NT-driven signaling cascade represents one of the most important challenges in modern neuropharmacology (Franco 2009).

In patients suffering from Parkinson’s disease, the progressive loss of dopaminergic neurons in the brain is compensated for by the administration of L-DOPA, a dopamine precursor. However, long term treatment with L-DOPA leads to numerous side effects, including the L-DOPA-induced dyskinesia (LID) (Pandey and Srivanitchapoom 2017). LID originates from an over sensitization of the Dopamine 1 dopamine receptor (DRD1), a Gs-protein GPCR, which leads to the pathological response to L-DOPA treatment (Santini et al. 2012). The use of an *ad hoc* developed pepducin would represent a novel treatment for LID, for it would allow modulating the activity of DRD1 while bypassing the malfunctioning receptor. In particular, acting upon the first intracellular step of the DRD1 signaling cascade, id est, the DRD1/Gαs-protein interaction, could allow for the modulation of the whole pathway. However, so far, no pepducins were ever developed for this purpose.

In this study, a lipidated cell-penetrating peptide mimicking the Gs-binding domain of the DRD1 was developed to target the DRD1/Gαs-protein interaction.

## Material and methods

### Design of the peptides

To develop a pepducin modulating the DRD1 signaling pathway, the DRD1/Gαs-protein interaction was targeted. A 2005 study by Kang et al. on the interaction between the intracellular loops of the human serotonin receptor type 6 (5-HT_6_, structurally and functionally similar to DRD1) and the α-subunit of the Gs protein, identified the interaction domain to be located on the C-terminal portion of the third intracellular loop (IL3) of 5-HT_6_, pinpointing two lysine residues (K262 and K265) to be crucial for the Gαs binding to the receptor (Kang et al. 2005). From a comparison between the sequences of the various Gs-coupled catecholamine receptors, the C-terminal portion of the IL3 resulted to be highly conserved (Fig.1). In particular Gαs-coupled receptor presented two lysine residues analogous of the aforementioned K262 and K265 of 5-HT_6_. Therefore, this portion of the receptor was used to generate the CPP/pepducins of interest.

**Fig.1.**
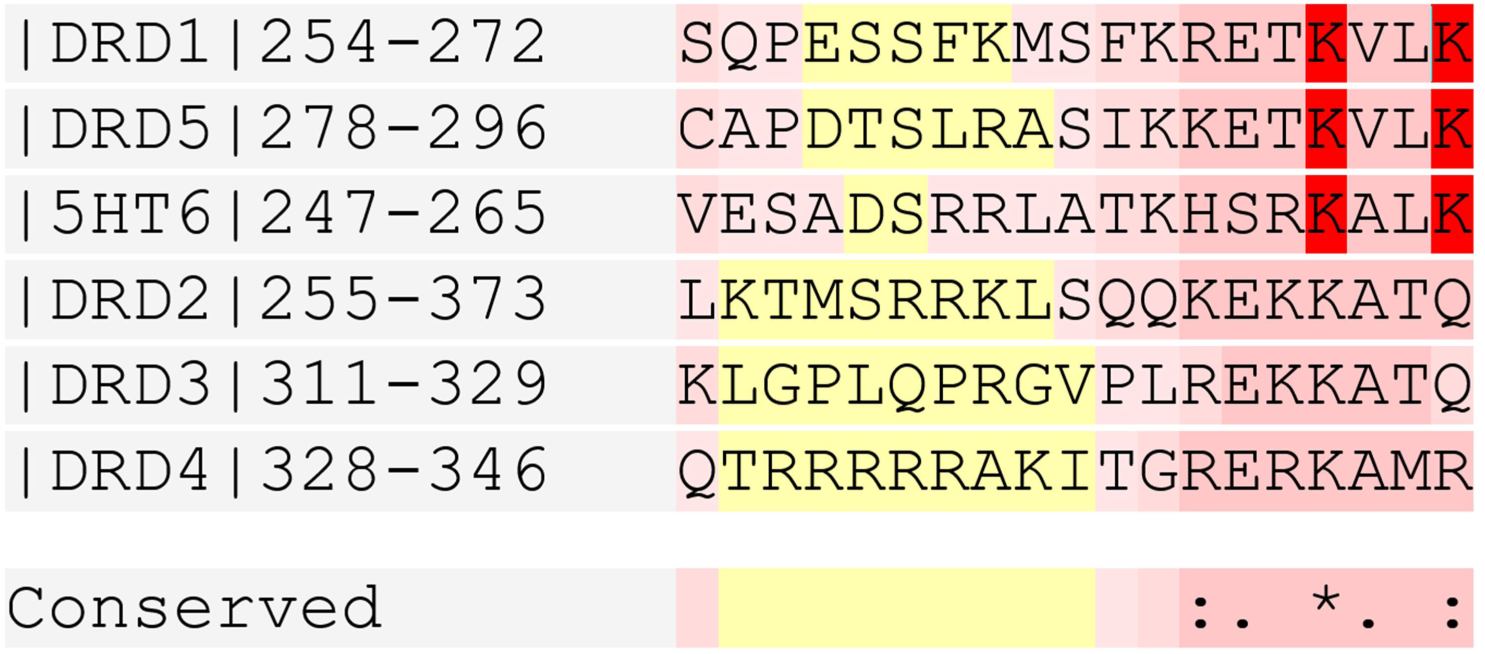
Sequence alignment of the C-terminal of the IL3 of dopaminergic and serotonergic receptors. DRD1, DRD5 and 5-HT6 are Gs-protein coupled receptors, whereas DRD2, DRD3 and DRD4 are Gi-coupled. The alignment shows highly conserved residues amongst receptors coupled to the same G-protein. In bright red: lysine residues crucial for Gαs subunit binding to the receptor as stated by Kang et al.,2005. Alignment performed via T-Coffee software (Notredame et al. 2000).

At first, computational estimation of the cell penetrating properties of the designed peptides was obtained using a CPP-prediction program: KELM-CPPpred. (Pandey et al. 2018) Sequences predicted to work as CPPs were then synthesized and tested as presented below. A lipid tail was added to the amino acid backbone to increase the likelihood of cell penetration (Lehto et al. 2017).

Peptides generated from the aforementioned portion of DRD1 were labeled as ‘D-1-R-Mo’, from ‘Dopamine 1 dopamine Receptor Modulators’ (later contracted to ‘Dyro’ for simplicity) followed by a serial number (Table 1). The fluorescent-probe-labeled version of a peptide that was employed during microscopy was labeled with an asterisk (*).

**Table 1.** Sequences of the peptides derived from the DRD1 IL3.

|  |  |
| --- | --- |
| Dyro 1 | Stearyl-KRETKVLK-NH<sub>2</sub> |
| Dyro 1* | Stearyl-KRETK(ε-5(6)carboxyfluorecein)VLK-NH<sub>2</sub> |
| Dyro 2 | KRETKVLK-NH<sub>2</sub> |
| Dyro 2* | KRETK(ε-5(6)carboxyfluorecein)VLK-NH<sub>2</sub> |
| Dyro 3 | Stearyl-PESSFKMSFKRETKVLKTLS-NH<sub>2</sub> |
| Dyro 4 | Palmitoyl-PESSFKMSFKRETKVLKTLS-NH<sub>2</sub> |
| Dyro 5 | RRRRRPESSFKMSFKRETKVLKTLS-NH<sub>2</sub> |
| Dyro 6 | PESSFKMSFKRETKVLKTLS-NH<sub>2</sub> |
| Dyro 7 | Stearyl-KMSFKRETKVLKTLS-NH<sub>2</sub> |

### Synthesis

The peptides were synthetized via Solid Phase Peptide Synthesis (SPPS) using Fluorenylmethyloxycarbonyl (Fmoc) chemistry protocols. The synthesis was performed either manually or by microwave-assisted SPPS (Biotage Alstra+, Biotage AB, Uppsala, Sweden) in dimethylformamide (DMF). H-Rink-Amide-ChemMatrix resin was used as solid support. Synthesis method and resin loading varied according to the sequence: due to the cumulative decrease in yield at every step of the synthesis, longer or sterically hindered peptides were synthetized manually on low-loading solid supports.

The carboxylic groups were activated with HCTU/HOBt (2-(6-Chloro-1H-benzotriazole-1-yl)-1,1,3,3-tetramethylaminium-hexafluorophosphate/6-Chloro-1-Hydroxy benzo-triazol; manual synthesis) or with DIC/Oxyma (N,N’-diisopropylcarbodiimide /Ethyl cyanohydroxyiminoacetate; microwave-assisted synthesis). Manual coupling of the amino acids was performed at room temperature. Couplings in microwave-assisted SPPS were performed at 70°C, although to avoid undesired side reactions, coupling of histidine, cysteine or arginine residues was performed at a maximum of 49°C. Coupling times ranged from 35min to 90min, depending on the position and nature of the coupled residue. Deprotection steps were performed in 20% piperidine. The fluorescent probe (5(6)-Carboxyfluorescein (FAM) was coupled to the side chain of the penultimate lysine residue from the N-terminus; for this purpose, a 4-Methyltrityl (Mtt)-protected lysine was integrated in the peptide, allowing for selective deprotection via 1%TFA/2.5%TIS/Dichloromethane cocktail.

The peptides were cleaved using a 95% Trifluoracetic acid (TFA)/2.5% water/2.5% triisopropylsilane cocktail for 3-4h and precipitated dropwise in ice-cold diethylether (Et_2_O). The obtained crude peptide was dried and then purified via high performance liquid chromatography (HPLC) on a C4, C8 or C18 column using a gradient of acetonitrile (ACN) / water containing 0.1% TFA. The purity and correct identity of the processed peptide products was assessed via analytical HPLC and by electrospray-time-of-flight mass spectrometery (ESI-TOF). After the purification and qualitative analysis, the peptides were lyophilized using a freeze-dryer, reconstituted in ultra-pure water and stored at -20°C before use.

### Cell lines

To assess whether the peptides had an effect on the DRD1-mediated signaling pathway, the SH-SY5Y cell line was employed, for, contrarily to the control cell line HeLa, SH-SY5Y cells express the aforementioned receptor. Furthermore, the neuroblastoma SH-SY5Y cell line is commonly used in PD research, whereas the adenocarcinoma HeLa cell line is widely employed in molecular biology research, assessing these cell lines as ideal model organisms for the purpose of this study (Landry et al. 2013; Xicoy et al. 2017). The cell line strains were already available in the laboratory, and the presence/absence of the receptor was assessed via real-time quantitative polymerase chain reaction (RT-qPCR) performed on cDNA obtained by retro-transcription of the total mRNA derived from the cell lines.

SH-SY5Y cells were grown in Minimum Essential Medium (MEM) (Sigma Aldrich Sweden), supplemented with 1x non-essential amino acids (NEAA), 1x L-Glutamate, 10% Fetal Bovine Serum (FBS), 200 µg/mL streptomycin and 200U/mL Penicillin (Invitrogen, Sweden). The medium was changed every 2 days, whereas cells were split 1 to 10 every 7 days.

HeLa cells were grown in Dulbecco’s Modified Eagle Medium (DMEM) (Sigma Aldrich Sweden), supplemented with 4.5g/L glucose, 1x NEAA, 1x L-Glutamate, 10% FBS, 200 µg/mL streptomycin and 200U/mL Penicillin (Invitrogen, Sweden). The medium was changed every 2 days, whereas cells were split 1 to 10 every 4 days.

### Epi-fluorescence microscopy

To establish the peptides function as CPPs, epi-fluorescence microscopy imaging was used. 15k cells were seeded in each well of a microscopy glass bottom black 96-well plate, left attaching overnight and then treated with various concentration of the fluorophore-conjugated peptide (ranging from 10nM to 50µM) at different points in time (1hr, 3hrs, 24hrs), followed by 15 min incubation with the nuclear stain (Hoechst 33342). To remove excess stain from the plate, the wells were then rinsed twice using Opti-MEM Reduced Serum Medium. To ensure cell viability, 200µL of the medium were added to each well before the microscopy.

Imaging was performed using a Leica DM/IRBE 2 epi-fluorescence microscope with 10X, 43X dry or 43X, 63 × 1.4 NA oil immersion objective. Samples were kept at 37°C at all times. Images were recorded via a Hamamatsu Orca-ER CCD camera. The system was controlled by Micro-Manager (Edelstein et al. 2014). Images were then processed using Image J (Rueden et al. 2017).

### Toxicity assay

The Pierce LDH Cytotoxicity Assay kit (Thermo Fisher) was used to assess the potential cytotoxic effects of the peptides due to the disruption of the membrane during cell penetration. The kit measures the quantity of lactate dehydrogenase (LDH, a cytosolic enzyme) present in the medium (Decker and Lohmann-Matthes 1988). Untreated cells were used as negative control, whereas Triton-X-treated-cells were used as positive control (for, being a detergent, it induces the complete disruption of the membrane and release of LDH in the medium). Cells were treated with different concentrations of the peptide (ranging from10nM to 100µM) at different time points (2hr to 24hrs). Data were plotted using GraphPad Prism 6.01 and normalized to the controls (Swift 1997).

### Dynamic light Scattering and ζ-potential

To determine the physical-chemical properties of the peptides and their propensity to particle formation, the hydrodynamic diameter and the ζ-potential of the peptides were measured using a Zetasizer Nano ZS (Malvern Instruments, United Kingdom). Measurements were performed at 37°C on 600µL of Opti-MEM containing various concentrations of the peptide (1µM, 5µM and 10µM) in Malvern Disposable Folded Capillary Zeta cells DTS1070 cuvettes. Opti-MEM was chosen for the characterization for it is the medium in which the cAMP experiments were performed.

### cAMP assay

DRD1 binds to the alpha subunit of a Gs-protein, which in turn activates the adenylyl cyclase (AC), increasing the production of cyclic-adenosine-monophosphate (cAMP), which is then employed as second messenger by the cell (Neve et al. 2004). To establish if cell treatment with the peptides led to variations in the levels of cAMP, indicating an effect on the DRD1 signaling pathway, the cAMP-Glo™ Max Assay kit (Promega) was employed. A clear bottom white 96-well plate was seeded with 15k cells/well. Half of the wells were treated with the peptide at various concentrations, half of the wells, instead, were treated with both the peptide at various concentrations and 0.25mM dopamine hydrochloride. This allowed to test whether the possible variations would occur only in presence of dopamine (hence only when the receptor was activated) or if the peptide was capable of interfering with the pathway in other ways. Untreated cells were used as negative control, whereas cholera toxin (capable of irreversibly activating the Gsα (Liu et al. 1992)) was used as positive control. The increase in cAMP levels derived from the activation of the DRD1-pathway after dopamine binding was measured on cells treated solely with 0.25mM of dopamine hydrochloride, a response-inducing concentration (Colombo et al. 2016) in an acceptable cytotoxic range (SOFIAN et al. 2014). All treatments were performed at different time points (1hr, 2hrs or 4hrs) in Opti-MEM Reduced Serum Medium supplied with IBMX and Ro 20-1724 (two cyclic nucleotide phosphodiesterase inhibitors, to prevent the degradation of the cAMP by native enzymes). The assay was then performed according to the manufacturers’ instructions. Data were plotted using GraphPad Prism and normalized to the controls (Swift 1997).

## Results

### Dyro 1 penetrates the cell membrane

Under Epi-fluorescence microscopy, Dyro 1* showed cell-penetrating properties already at 1µM after 1hr of treatment (Fig.2). Dyro 2*, the unstearylated form of Dyro 1, instead, was not capable of crossing the cell membrane, suggesting that the lipid tail is implicated in the translocation mechanism.

**Fig. 2.**
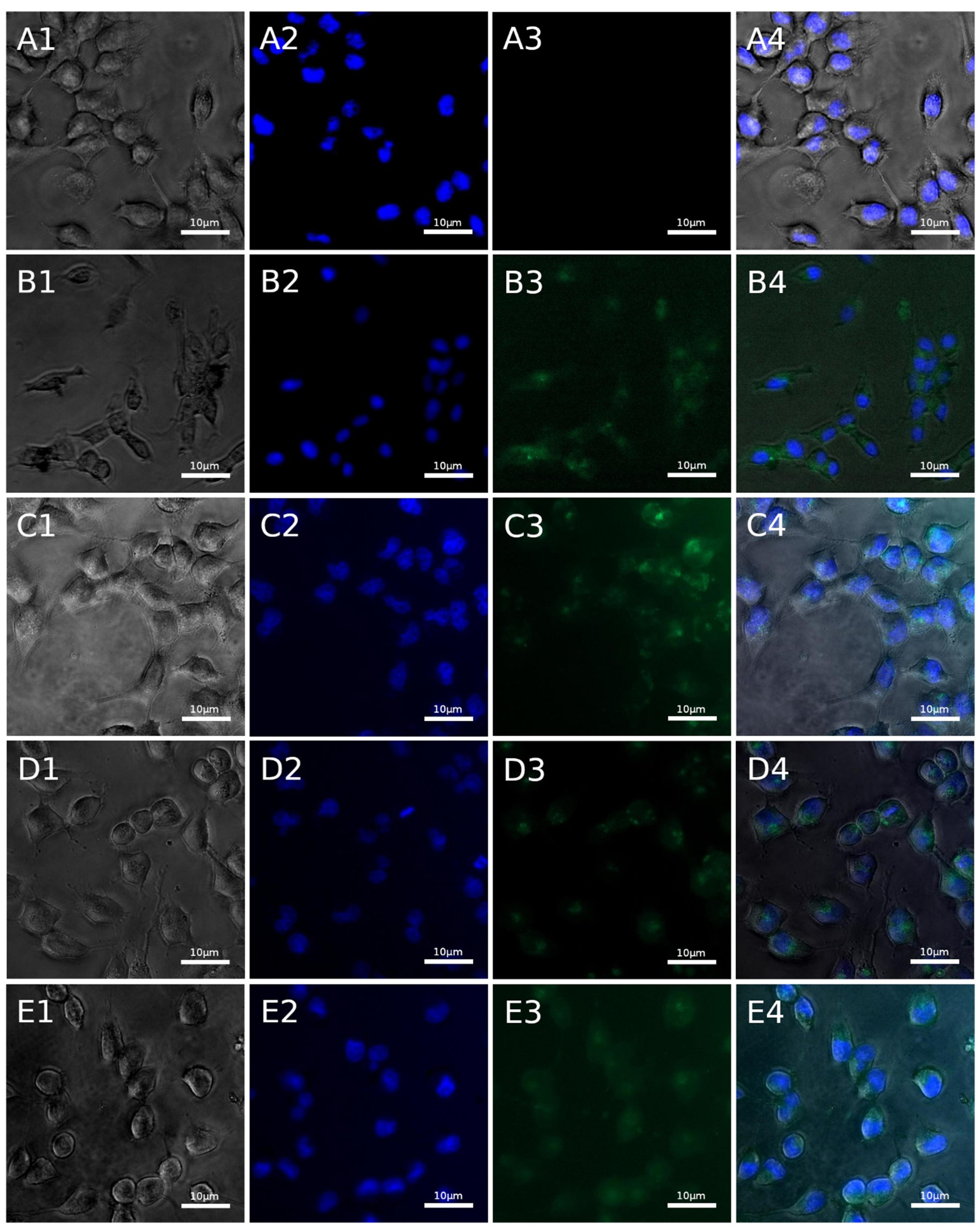
Dyro 1* on HeLa cells. Columns from left to right: Transmission channel, Hoechst channel (nuclear stain), fluorescein channel, merged channels. Rows: A) Cell treated with 1µM carboxyfluorescein solution for 1h (Control); B) Cells treated with 1µM Dyro 1* for 1h; C) Cells treated with 10µM Dyro 1*, 1h treatment; D) 1µM Dyro 1*, 24hrs treatment; E) 10µM Dyro 1*; 24hrs treatment.

Stearylated peptides derived from a larger portion of the DRD1-IL3 (from K261-S275 onwards) were not soluble in water and were therefore discarded. Substitution of the lipid tail with a polyarginine tail (a known CPP (Fuchs and Raines 2005)) did not allow for solubilization in water either, indicating that the problem originates from the sequence itself.

### Assessing the working concentration

Utilizing the aforementioned Pierce LDH Cytotoxicity Assay, the concentration of Dyro 1 sufficient to induce rupturing of the membrane in half of the cells treated (EC50) was determined to be 6.72µM after 24hrs (Fig.3) and 100µM after 2hrs (Supplementary figure S.1). Concentrations that induced less than 20% of membrane rupture were deemed to be acceptable working concentrations for that or shorter treatment times.

**Fig. 3.**
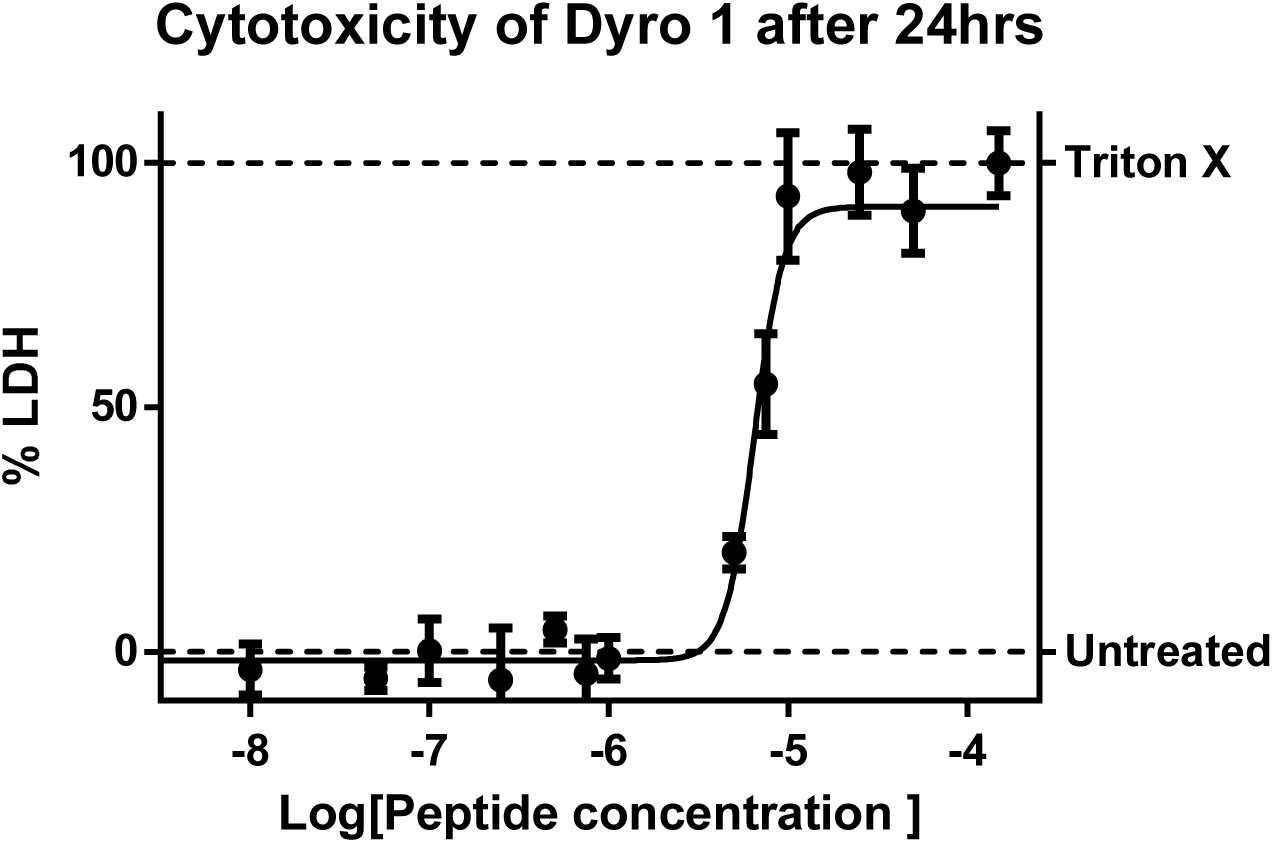
Cytotoxicity of Dyro 1 on SH-SY5Y cells after 24hrs treatment. SH-SY5Y cells were treated with increasing concentrations of Dyro 1 for 24hrs. The quantity of lactate dehydrogenase (LDH, a cytosolic enzyme) present in the medium was measured as an indicator for membrane disruption and cell death. Data were normalized to the controls: untreated cells were used as negative control (physiological release of LDH in the cell medium; 0%), whereas cells treated with Triton X were used as the positive control (being a detergent, Triton X induces membrane disruption and complete LDH release; 100%). Control samples were examined via microscopy before LDH measurements to check for viability. The experiment was performed three times in quadruplicates (p value = 0.0012). Calculated EC50 = 6.72µM.

### Dyro 1 aggregates in particles

DLS and ζ-potential measurements were performed to characterize Dyro 1 in solution, as stated in the Material and methods section. The hydrodynamic diameter measurements revealed that Dyro 1 forms particles, whose size increases exponentially depending on the concentration (Supplementary figure S2). The formation of particles was confirmed also by the ζ-potential, which assessed the peptide to be in rapid coagulation range at the concentrations of interest (Maha et al. 2016). This suggests that hydrophobic interactions between the lipid tails of Dyro 1 molecules led to the formation of particles. Given that numerous CPP models rely on the interaction between lipopeptidic aggregates and the cell membrane to explain the mechanism of cell penetration, these data can provide an explanation on why the unstearylated versions of the peptide proved not to be capable to enter the cell.

### Dyro 1 selectively increases cAMP levels in cells expressing DRD1

To study the possible effects of Dyro 1 on the dopamine signaling pathway, variations in cAMP levels were measured after treatment. Experiments were performed as stated in Material and methods. SH-SY5Y cells (expressing DRD1) showed a concentration-dependent increase in cAMP levels after treatment with the peptide (Supplementary figure S3). In particular, a 4hrs treatment was sufficient to induce an up to 37% increase in cAMP levels compared to the controls (untreated cells and cholera toxin treated cells; Fig.4). This is almost a 2-fold increase compared to the 23.2% increase in cAMP levels induced by DRD1 activation via dopamine-binding measured by treating cells solely with 0.25mM dopamine hydrochloride for 4hrs and comparing the data to the aforementioned controls (Fig.4). This effect was not observed in the control cell line (HeLa cells, not expressing DRD1), which showed no significant increase in cAMP levels after 4hrs of treatment with Dyro 1 (Supplementary figure S.4), suggesting that the presence of the receptor plays a role in the peptide-induced cAMP production.

**Fig. 4.**
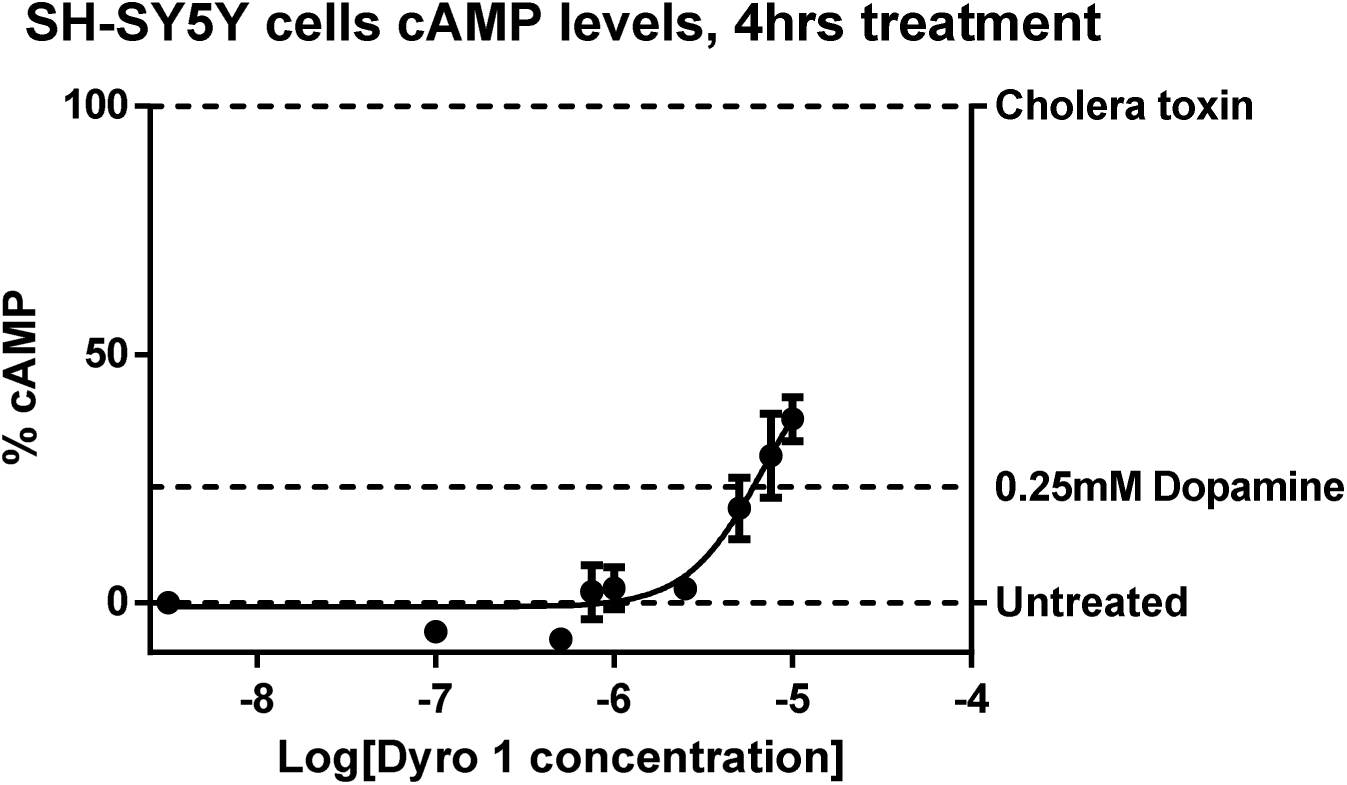
cAMP levels of SH-SY5Y cells after 4hrs treatment. SH-SH5Y cells (expressing DRD1) were treated with increasing concentrations of Dyro 1 for 4hrs. In order not to incur in the peptide cytotoxic effect, 10 µM Dyro 1 was the maximum peptide concentration used. Data were normalized to the controls: untreated cells (0%) and cholera-toxin-treated cells (100%; cholera toxin is known to irreversibly activate AC, thereby setting the maximal cAMP production levels). Cells treated solely with 0.25mM dopamine hydrochloride were employed to measure the variation of cAMP levels following DRD1 response to ligand binding (23.2% in the graph). Samples were examined via microscopy before cAMP level measurements to check for viability. The experiment was performed four times in quintuplicates for statistical significance (use of one-tail multiple T-tests).

Furthermore, SH-SH5Y cells that were co-treated for 4hrs with different concentrations of Dyro 1 and 0.25mM dopamine hydrochloride, showed a constant ≈25% concentration-independent increase in cAMP levels (Supplementary figure S5) similar to the +23.3% cAMP effect observed in the cells that had been treated solely with dopamine. This effect was therefore attributable to the presence of dopamine in the treatment mixture and not to the peptide.

## Discussion and conclusions

Parkinson’s disease is, characterized by the progressive loss of dopaminergic neurons in the periphery and in the central part of the nervous system. Patients suffering from this disorder are commonly treated with L-DOPA, which is capable of temporarily compensating for the loss of endogenous dopamine. However, long-term L-DOPA treatment overcharges the remaining dopaminergic receptors, leading to their over sensitization and the consequential onset of numerous side-effects. In particular L-DOPA-induced-dyskinesia (LID) originates from the over sensitization of the dopamine 1 receptor (DRD1) (Santini et al. 2012). Bypassing the malfunctioning receptors by targeting directly the dopaminergic pathways from the inside of the cell might represent a novel approach to contrast the occurrence of these side effects. Pepducins, i.e. lipidated cell-penetrating peptides designed to mimic the active site of a protein of interest, are good candidates for this purpose.

In this study, stearylated peptides derived from the intracellular active site of the dopamine 1 receptor, were designed to target the DRD1/Gαs-protein interaction. Following previous studies on pepducins targeting the GPCR/Gαs interaction, the portion of interest was individuated in the third intracellular loop (IL3) of DRD1 (O’Callaghan et al. 2012). Furthermore, from a comparison between the sequences of various Gs-coupled catecholamine receptors (Fig.1), the last portion of the IL3 resulted to be the most conserved. In particular, two lysine residues (K269 and K272) were referable to the K262 and K265 residues of the IL3 of serotonin receptor type 6, which are known to be crucial in the receptor/G-protein interaction (Kang et al. 2005). Dyro 1, the C-terminal stearylated peptide derived from the K265-K272 portion of the DRD1-IL3, proved to be the best candidate.

Under epi-fluorescent microscopy, the fluorescein-labeled Dyro 1* was assessed work as a cell-penetrating peptide. However, its unstrearylated version proved not to be capable of autonomously crossing the cell membrane, suggesting that the lipid tail plays a crucial role in the translocation mechanism. This is consistent not only with previous studies on CPPs, where lipidation is often used to increase the efficiency of the penetration (Lehto et al. 2017), but on pepducins in general, where the presence of the tail is a conditio sine qua non the peptide can work as a CPP on its own (Zhang et al. 2015). Furthermore, from the DLS analysis and ζ-potential measurements, Dyro 1 proved to be capable to form particles in solution. This, together with the necessity for the lipid tail to enter the cell, suggested that the translocation mechanism relies on hydrophobic interactions between the lipopeptidic particles and the cell membrane, in line with the most common CPP penetration mechanism hypothesis (Bechara and Sagan 2013).

After treatment, Dyro 1 selectively induced a concentration-dependent increase in cAMP levels in cells expressing DRD1, while presenting no effect on the control cell line, which did not express the receptor. This indicated that the presence of the receptor was necessary to induce the cellular response to treatment, which is in line with previous in vivo studies on pepducins activity performed on knock-out mice (Kaneider et al. 2007; Sevigny et al. 2011). This increase is comparable to the one observed in a 2014 study by Carr et al., where two pepducins derived from the IL3 of the beta-adrenergic 2 receptor (a Gαs-coupled GPCR) were capable of inducing an up to +40% concentration-dependent effect on cAMP levels in cells expressing the receptor (Carr et al. 2014).

Furthermore, the Dyro 1 seemingly showed no effect on cells that had been co-treated with dopamine hydrochloride, an agonist of DRD1, suggesting that the peptide does not have an effect on an already activated signaling pathway. This could be explained in the light of the mechanism of action of the peptide: pepducins have been found to be capable of mimicking and/or inducing the intracellular on-conformation of the receptor of interest (O’Callaghan et al. 2012). An already-active receptor would therefore not be affected by the action of these peptides.

In conclusion, experiments on Blood Brain Barrier (BBB) models might be used to assess the ability of Dyro 1 to enter the central nervous system. If successful, mice models could be employed to study its effects on the brain and in particular if it could be used as a novel drug or co-treatment for patients suffering from PD and LID which require the DRD1-signally pathway to be artificially activated, but where the receptor is not properly functioning and the use of extracellular ligands has proved to induce major side effects. In particular, the peptide might be employed to substitute partially, or, in combination with similar peptides, completely, L-DOPA in the treatment of Parkinson’s disease.

## Authors contribution

NL conceived the study, ÜL supervised the study, NL designed and performed the experiments, MG participated in the experiments, NL wrote and revised the manuscript, MG and ÜL contributed to revise the manuscript. All authors have read and approved the final manuscript.

