## Supplementary figures for "Mimicry of dopamine 1 receptor signaling with Cell-Penetrating Peptides"

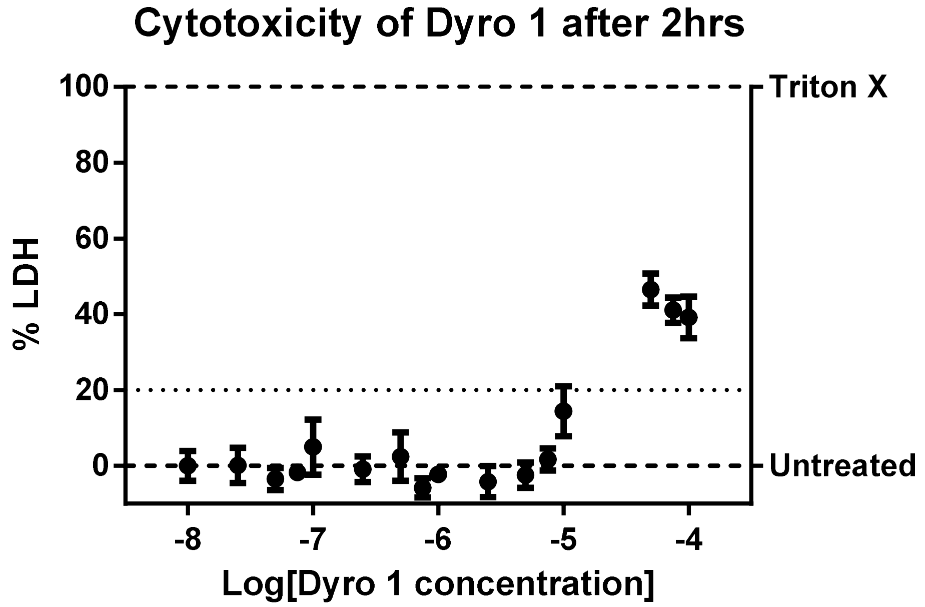


**Supplementary figure S1 – Cytotoxicity of Dyro 1 on SH-SY5Y cells after 2hrs treatment**. SH-SY5Y cells were treated with increasing concentrations of Dyro 1 for 24hrs. The quantity of lactate dehydrogenase (LDH, a cytosolic enzyme) present in the media was measured as an indicator for membrane disruption and cell death. Data were normalized to the controls: untreated cells were used as negative control (physiological release of LDH in the cell medium; 0%), whereas cells treated with Triton X were used as the positive control (being a detergent, Triton X induces membrane disruption and complete LDH release; 100%). Control samples were examined via microscopy before LDH measurements to check for viability. The experiment was performed three times in triplicates. Due to experimental necessities, the examined Dyro 1 concentration range was 10nM - 100µM, which was not enough to build a curve. However, comparing the data to the controls, it was still possible to determine the working concentration range (being its maximum, the peptide concentration inducing 20% of membrane rupture at that timepoint). The EC50 was experimentally determined to be at 100µM.

| | Particle size (d.nm) | ζ-potential (mV) |
|---|---|---|
| 1 µM | 143,5 | -5,00 |
| 5 µM | 366,7 | -3,63 |
| 10 µM | 1521 | +2,97 |

**Supplementary figure S2 – Dyro 1 particles characterization via DLS.** Particle size (diameter expressed in nm) and ζ-potential (expressed in mV) and were measured using a Zetasizer Nano ZS. Particle size measurements shows a concentration-dependent increase in size, suggesting the formation of particles. ζ-potential measurements set Dyro 1 to be in the rapid coagulation range.

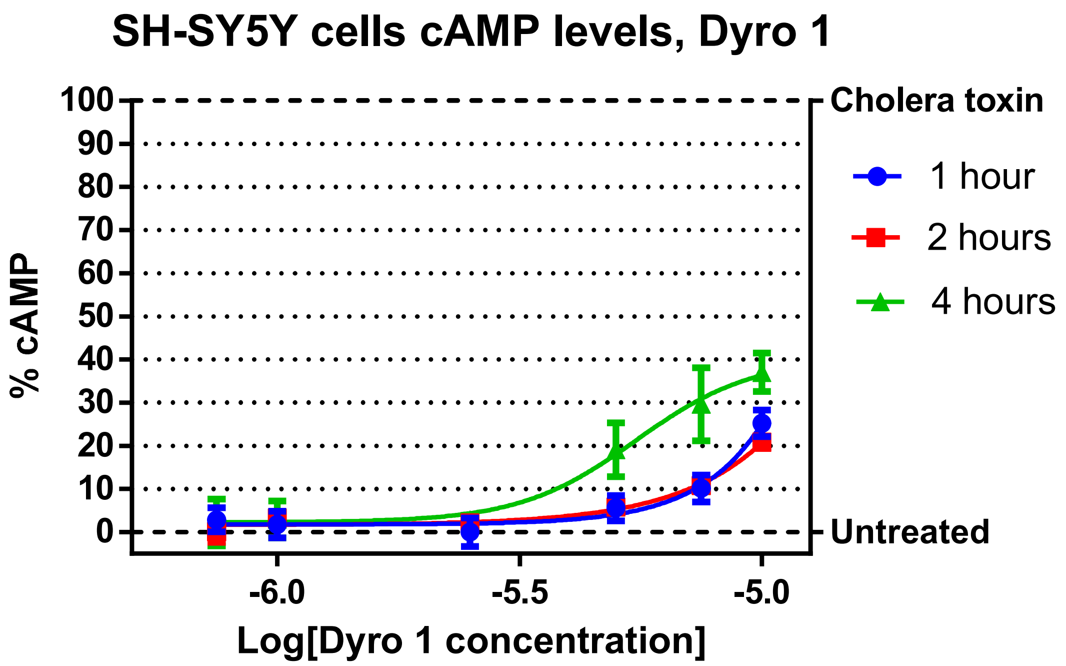


**Supplementary figure S3 – cAMP levels of SH-SY5Y cells after Dyro 1 at different treatment times**. SH-SH5Y cells (expressing DRD1) were treated with increasing concentrations of Dyro 1 for 1h, 2hrs or 4hrs. In order not incur in the peptide cytotoxic effect, 10 µM Dyro 1 was the maximum peptide concentration used. Data were normalized to the controls: untreated cells (0%) and cholera-toxin-treated cells (100%). Controls were performed for each set of samples. Samples were examined via microscopy before cAMP level measurements to check for viability. The 1h and 2hrs measurements were performed twice in triplicates (confirmed by T-test, α=0.05). The 4hrs measurements were performed four times in quintuplicates for statistical significance (confirmed by multiple one tailed multiple T-tests, α=0.05).

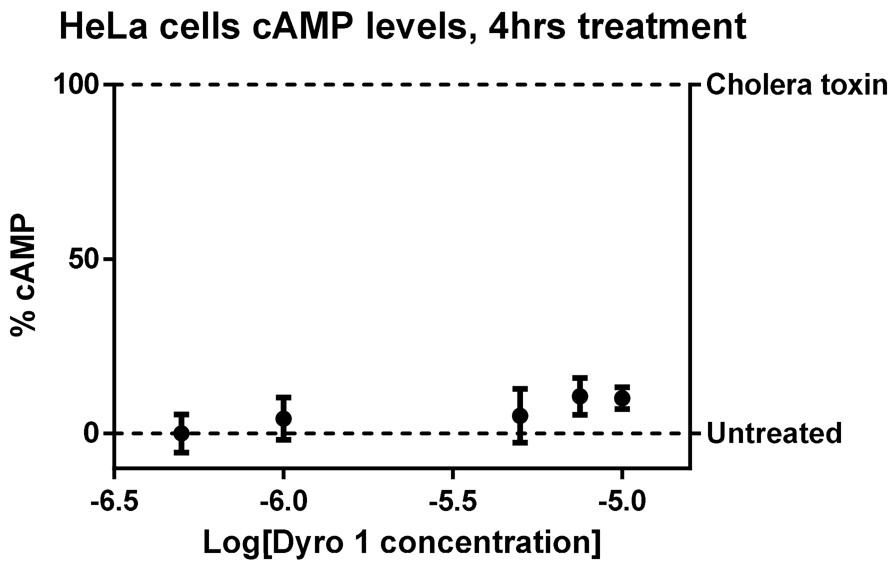


**Supplementary figure S4 – cAMP levels of HeLa cells after 4hrs treatment**. HeLa cells (not expressing DRD1) were treated with increasing concentrations of Dyro 1. Data were normalized to the controls: untreated cells (0%) and cholera-toxin-treated cells (100%). Experiment was performed three times in triplicates (p value < 0.0001, one-tailed analysis).

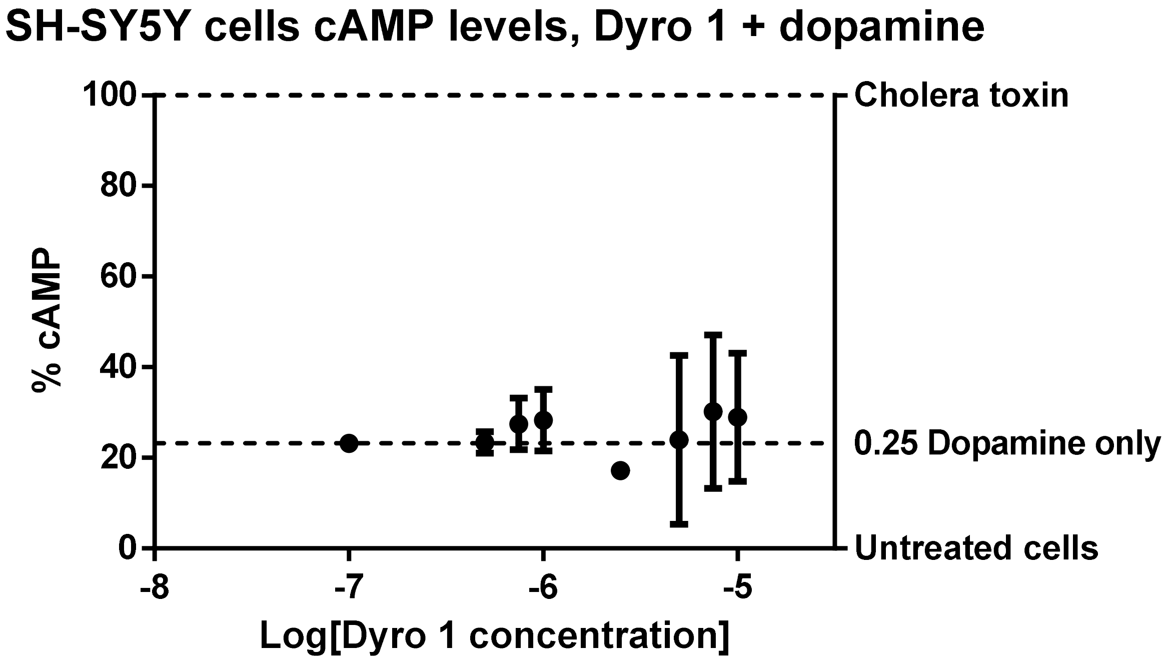


**Supplementary figure S5 – cAMP levels of SH-SY5Y cells after 4hrs treatment**. SH-SH5Y cells (expressing DRD1) were treated with increasing concentrations of Dyro 1 and 0.25mM dopamine hydrochloride for 4hrs. In order not incur in the peptide cytotoxic effect, 10 μM Dyro 1 was the maximum peptide concentration used. Data were normalized to the controls: untreated cells (0%) and cholera-toxin-treated cells (100%). Data obtained by cells treated solely with 0.25mM dopamine hydrochloride is reported in the graph. Samples were examined via microscopy before cAMP level measurements to check for viability. The experiment was performed four times in quintuplicates for statistical significance.
